## Supporting Figures for "Structure and dynamics of a nanodisc by integrating NMR, SAXS and SANS experiments with molecular dynamics simulations"

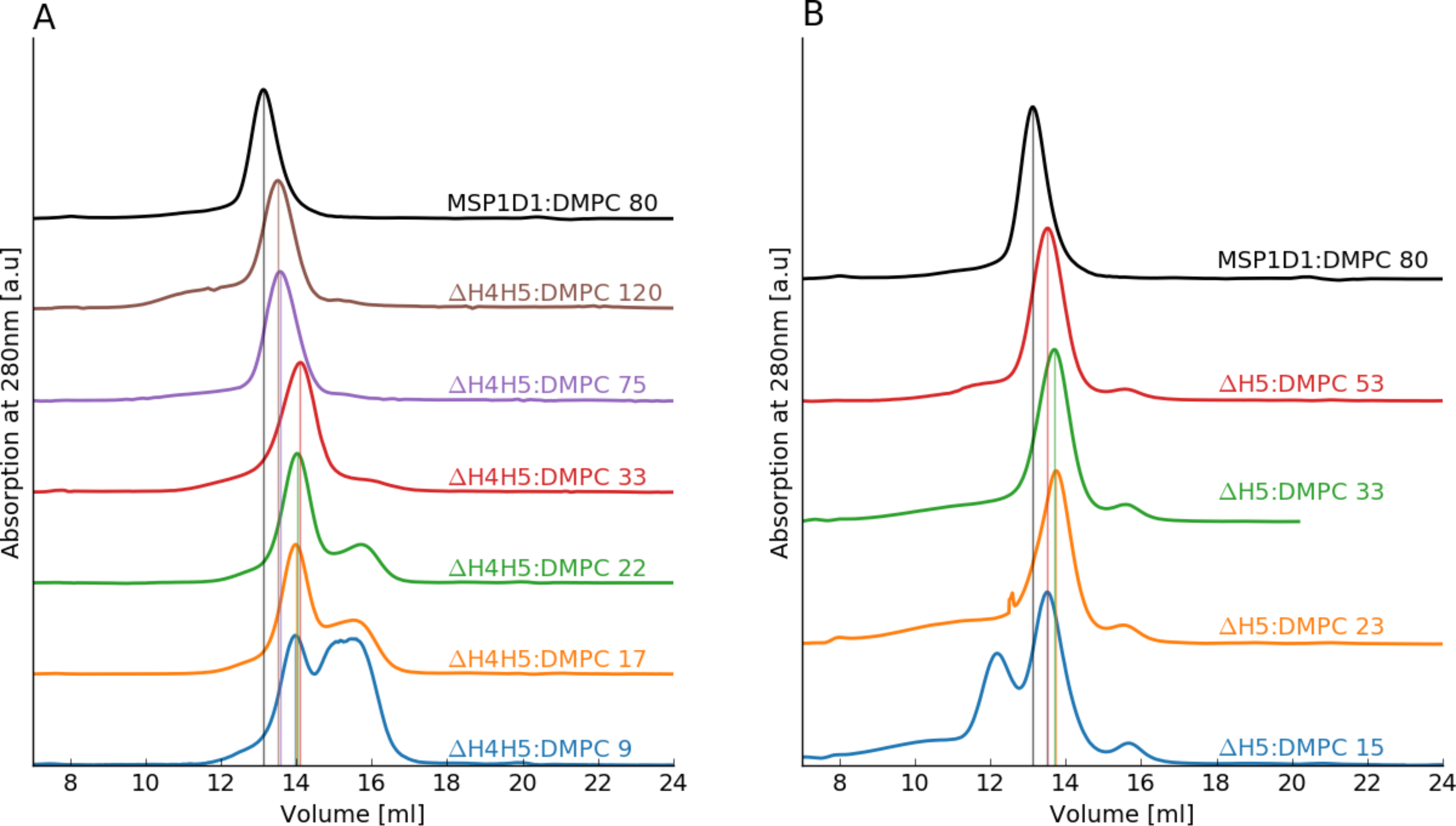
A
MSP1D1:DMPC 80
ΔH4H5:DMPC 120
ΔH4H5:DMPC 75
ΔH4H5:DMPC 33
ΔH4H5:DMPC 22
ΔH4H5:DMPC 17
ΔH4H5:DMPC 9
Absorption at 280nm [a.u]
Volume [ml]
8
10
12
14
16
18
20
22
24
B
MSP1D1:DMPC 80
ΔH5:DMPC 53
ΔH5:DMPC 33
ΔH5:DMPC 23
ΔH5:DMPC 15
Absorption at 280nm [a.u]
Volume [ml]
8
10
12
14
16
18
20
22
24

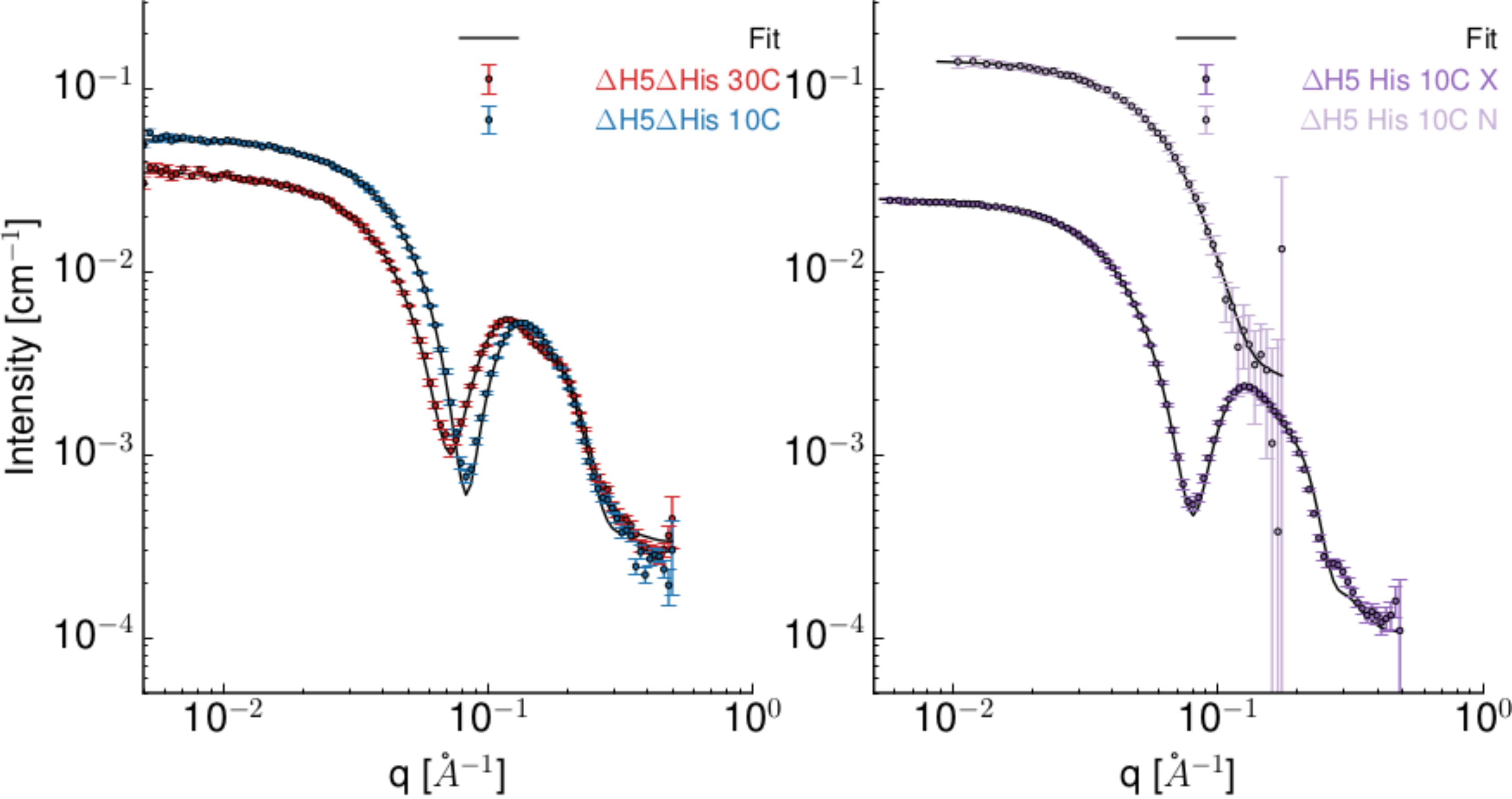


| | Static ΔH5ΔHis 30C | Static ΔH5ΔHis 10C | SEC-SAS ΔH5 His 10C |
|---|---|---|---|
| $\chi^2_{reduced}$ | 2.40 | 3.76 | 1.95 |
| | **Fitting Parameters** | | |
| Axis Ratio | 1.3±0.1 | 1.4±0.1 | 1.3±0.4 |
| $A_{Head}$ | 60± 3 Å$^2$ | 52± 2 | 55± 5 |
| $H_{Belt}$ | 24* Å$^2$ | 24* Å$^2$ | 24* Å$^2$ |
| $N_{Lipid}$ | 104± 9 | 102± 7 | 65±13 |
| $CV_{belt}$ | 0.97±0.02 | 1* | 1* |
| $CV_{lipid}$ | 1.044±0.007 | 1.003±0.007 | 1±0.02 |
| $Back_{neutron}$ | 0± 0 | 0± 0 | -0.0036±0.006 |
| $Scale_{x-ray}$ | 1.2±0.2 | 1.2±0.1 | 1.1±0.3 |
| $Scale_{neutron}$ | - | - | 1.7±0.5 |
| | **Results From Fits** | | |
| $H_{lipid}$ | 38 Å | 41 Å | 40 Å |
| $H_{tails}$ | 26 Å | 29 Å | 28 Å |
| $R_{major}$ | 36 Å | 34 Å | 27 Å |
| $R_{minor}$ | 28 Å | 25 Å | 21 Å |
| $W_{belt}$ | 9 Å | 9 Å | 10 Å |

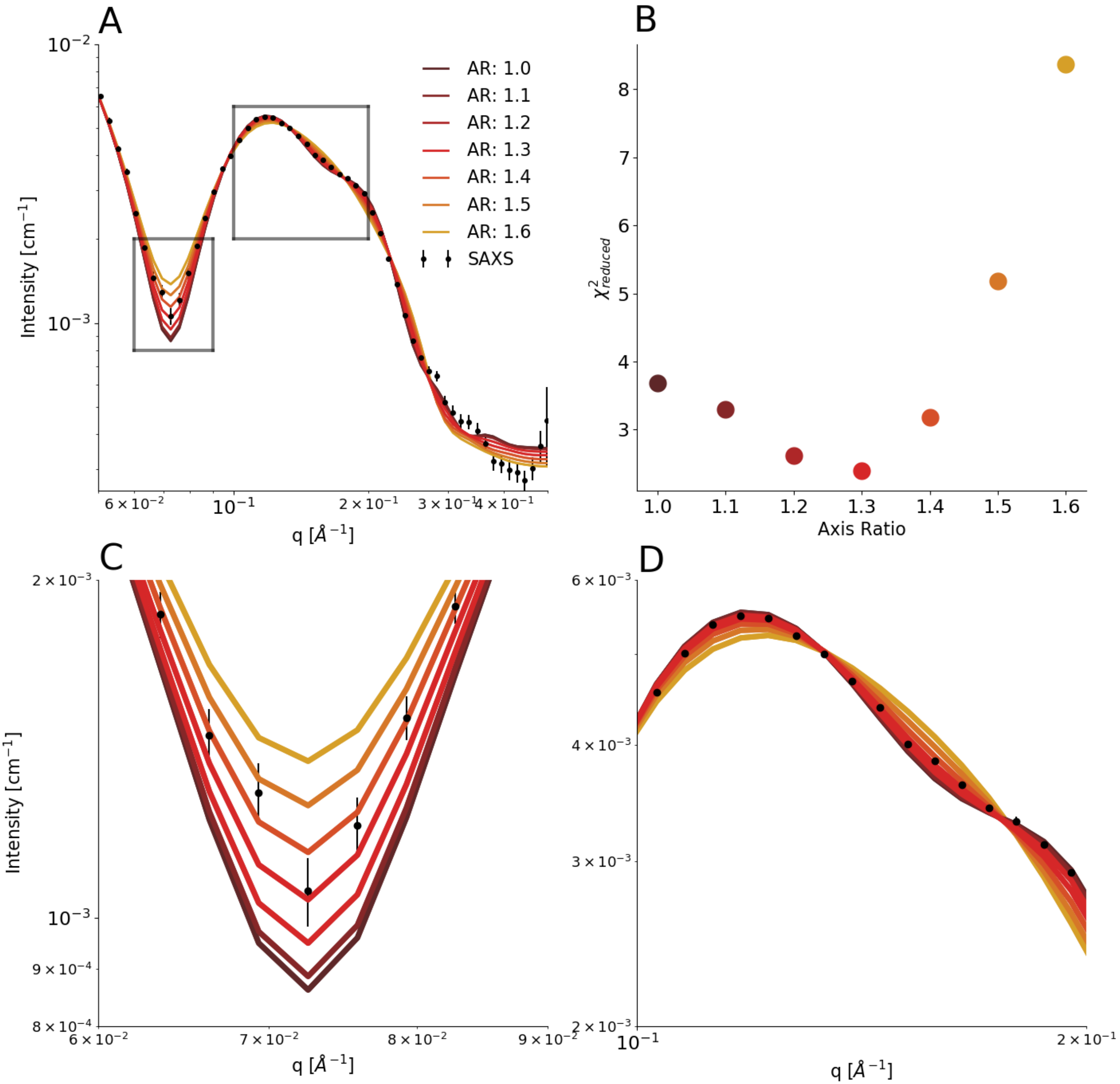

A
B
C
D
AR: 1.0
AR: 1.1
AR: 1.2
AR: 1.3
AR: 1.4
AR: 1.5
AR: 1.6
SAXS
Intensity [cm$^{-1}$]
q [Å$^{-1}$]
$\chi^2_{reduced}$
Axis Ratio

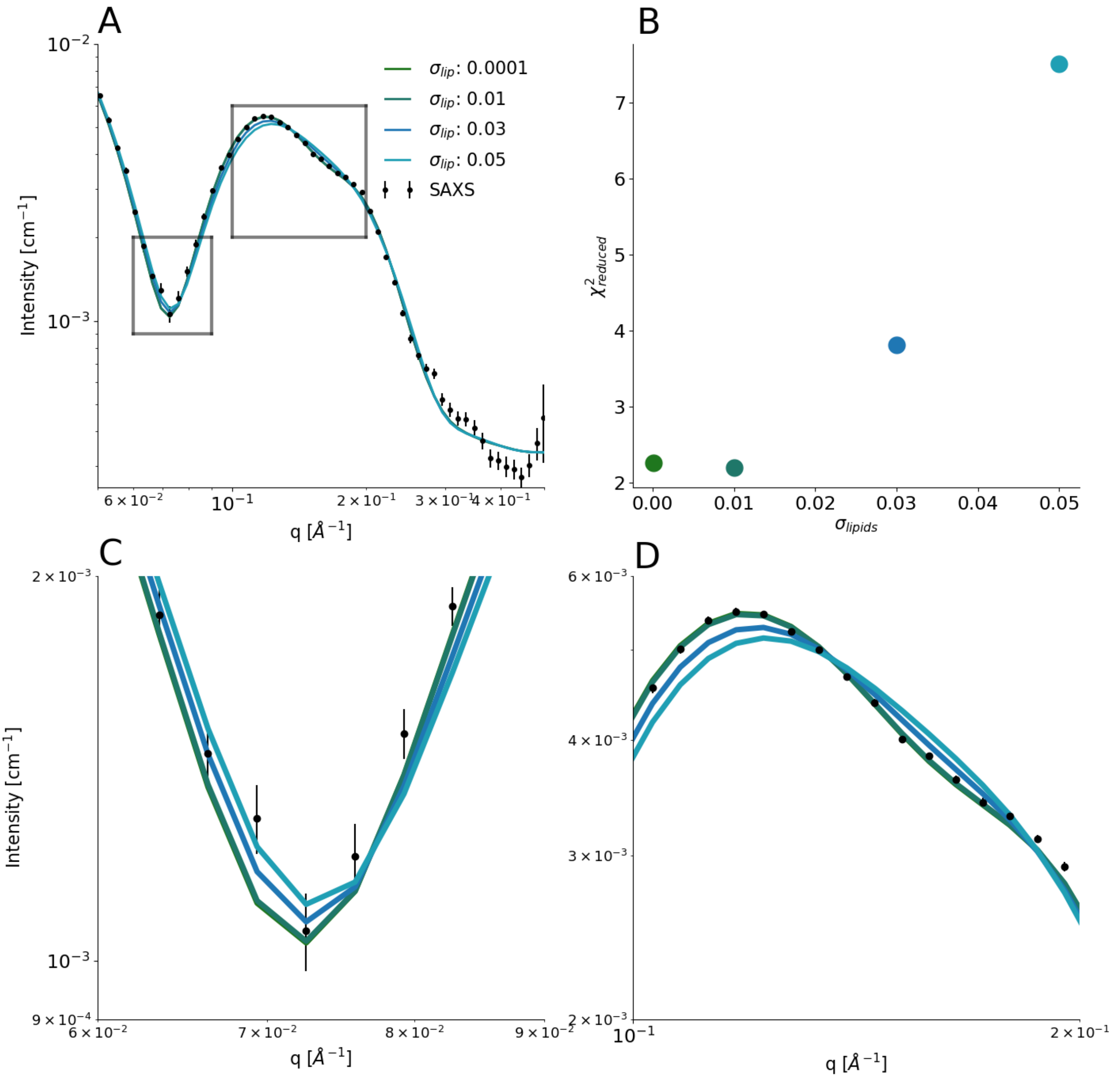
A
B
C
D
$\sigma_{lip}$: 0.0001
$\sigma_{lip}$: 0.01
$\sigma_{lip}$: 0.03
$\sigma_{lip}$: 0.05
SAXS
Intensity [cm$^{-1}$]
q [Å$^{-1}$]
$\chi^2_{reduced}$
$\sigma_{lipids}$

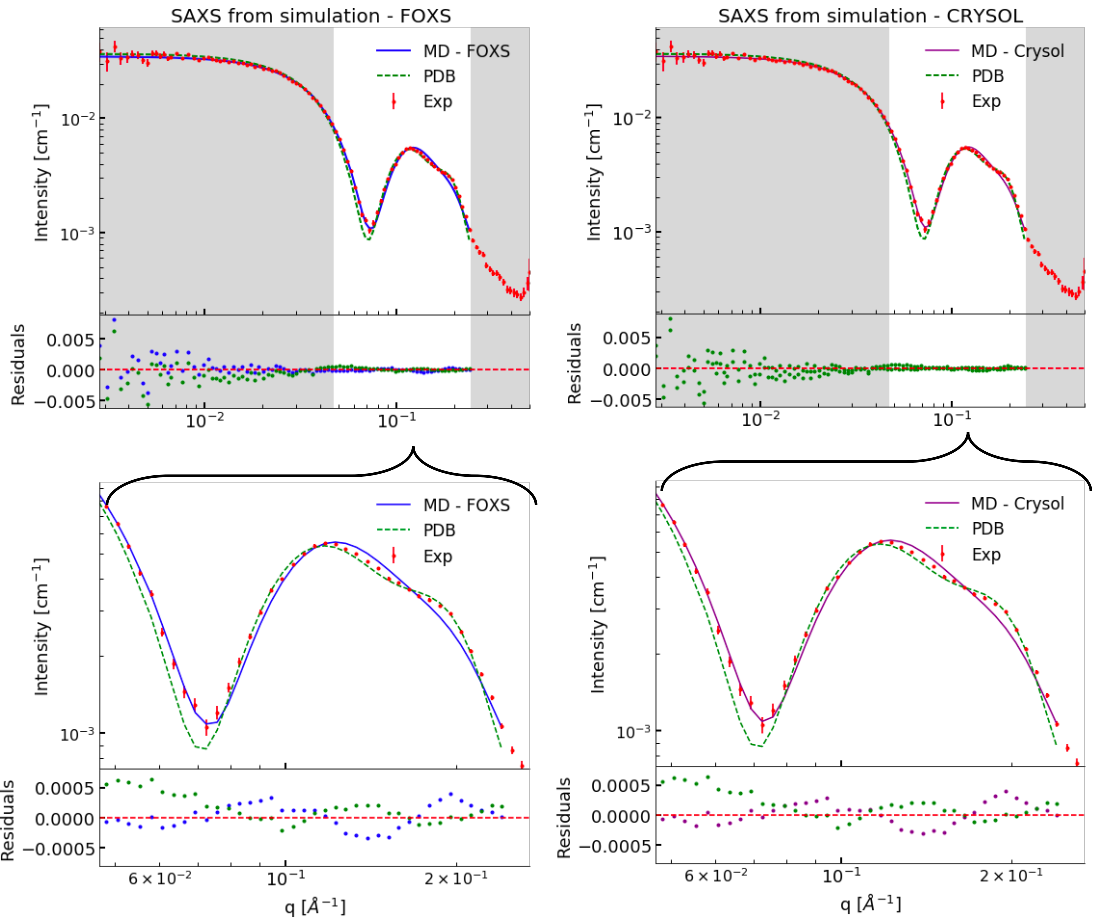

SAXS from simulation - FOXS
SAXS from simulation - CRYSOL
MD - FOXS
MD - Crysol
PDB
Exp
Intensity [cm$^{-1}$]
Residuals
q [$Å^{-1}$]
$10^{-2}$
$10^{-3}$
$10^{-1}$
0.005
0.000
−0.005
0.0005
0.0000
−0.0005
$6\times10^{-2}$
$2\times10^{-1}$

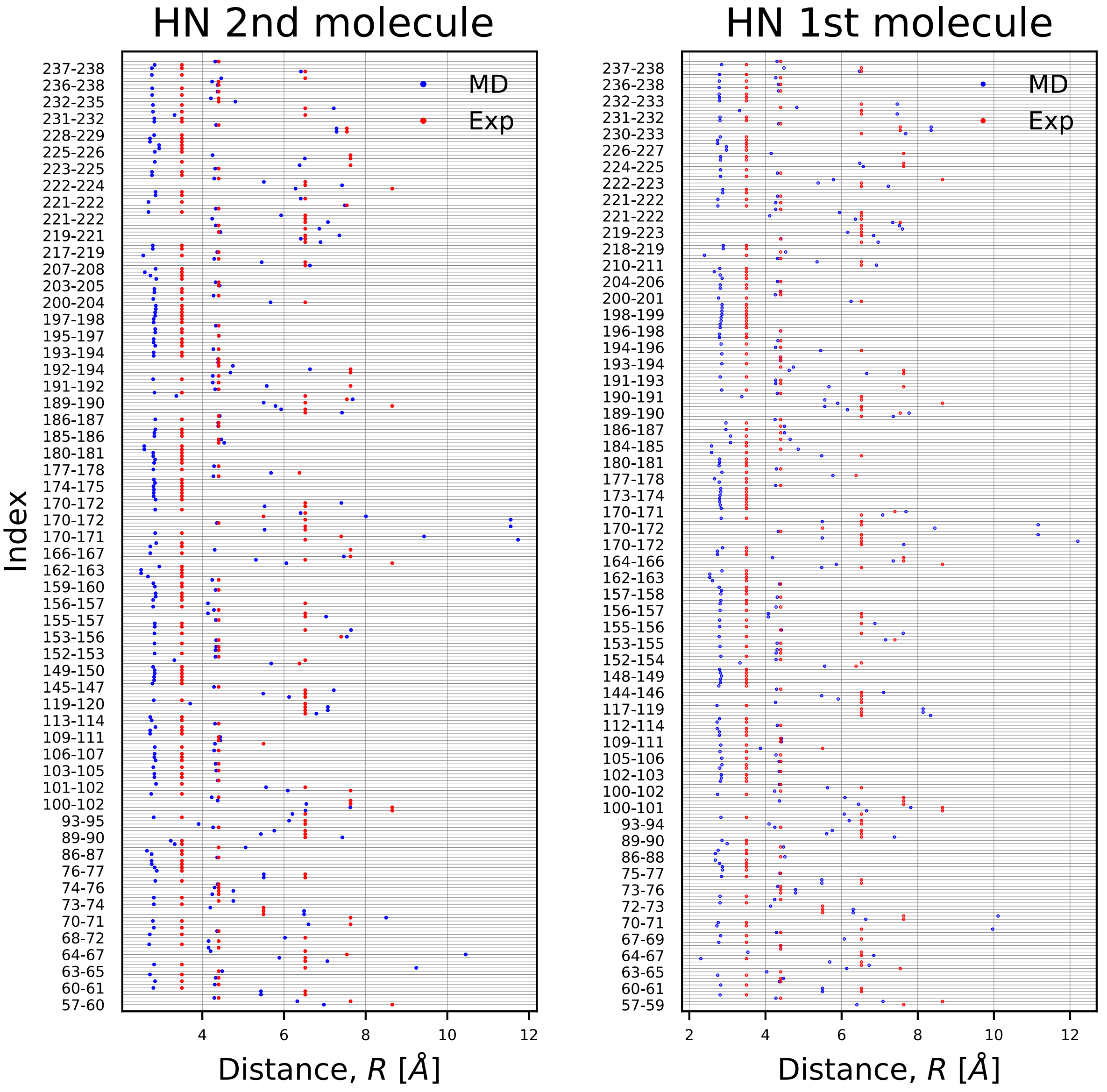

HN 2nd molecule
HN 1st molecule
Index
MD
Exp
Distance, $R$ [$Å$]
237-238
236-238
232-235
231-232
228-229
225-226
223-225
222-224
221-222
221-222
219-221
217-219
207-208
203-205
200-204
197-198
195-197
193-194
192-194
191-192
189-190
186-187
185-186
180-181
177-178
174-175
170-172
170-172
170-171
166-167
162-163
159-160
156-157
155-157
153-156
152-153
149-150
145-147
119-120
113-114
109-111
106-107
103-105
101-102
100-102
93-95
89-90
86-87
76-77
74-76
73-74
70-71
68-72
64-67
63-65
60-61
57-60
4
6
8
10
12
237-238
236-238
232-233
231-232
230-233
226-227
224-225
222-223
221-222
221-222
219-223
218-219
210-211
204-206
200-201
198-199
196-198
194-196
193-194
191-193
190-191
189-190
186-187
184-185
180-181
177-178
173-174
170-171
170-172
170-172
164-166
162-163
157-158
156-157
155-156
153-155
152-154
148-149
144-146
117-119
112-114
109-111
105-106
102-103
100-102
100-101
93-94
89-90
86-88
75-77
73-76
72-73
70-71
67-69
64-67
63-65
60-61
57-59
2
4
6
8
10
12

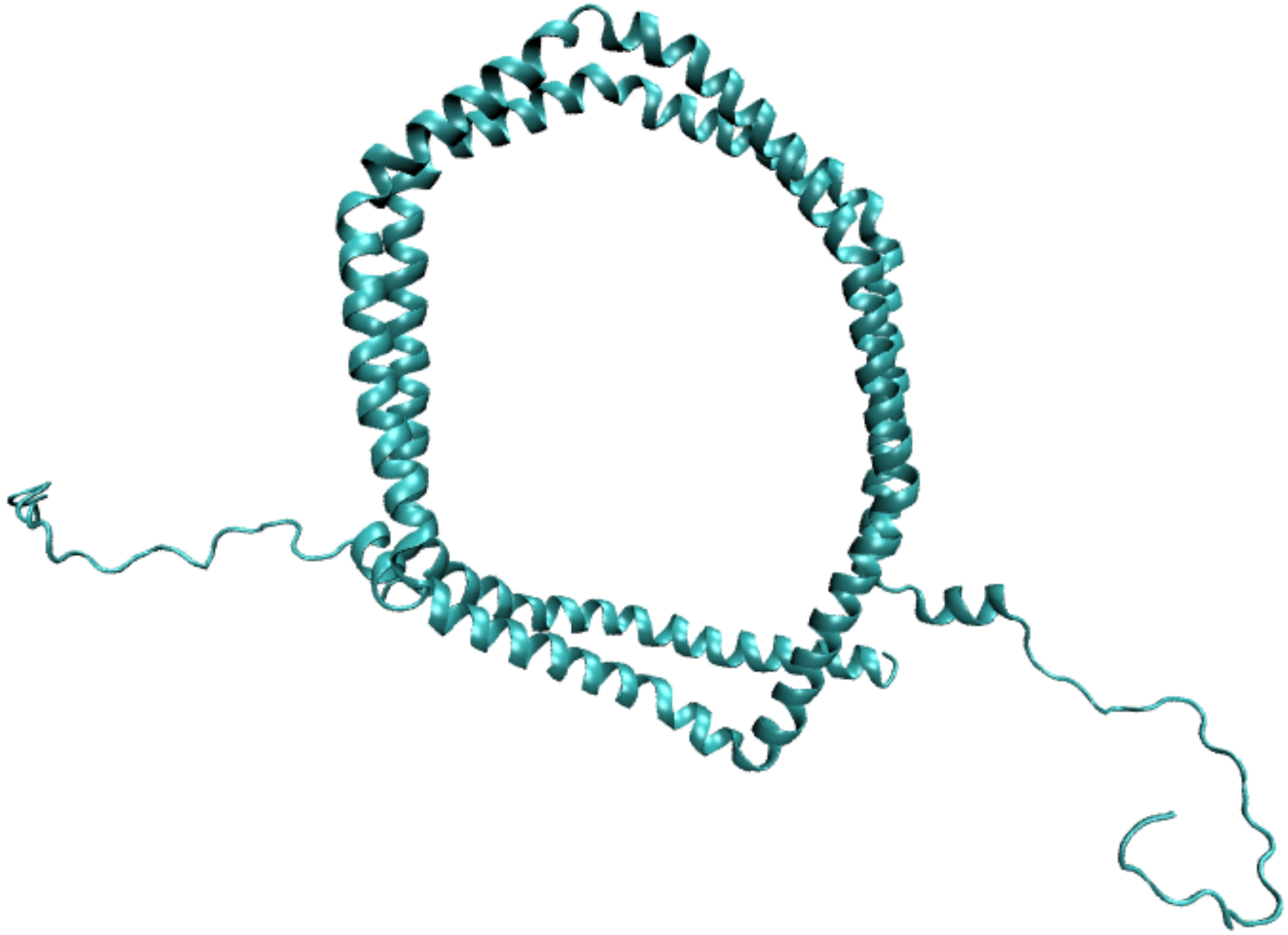

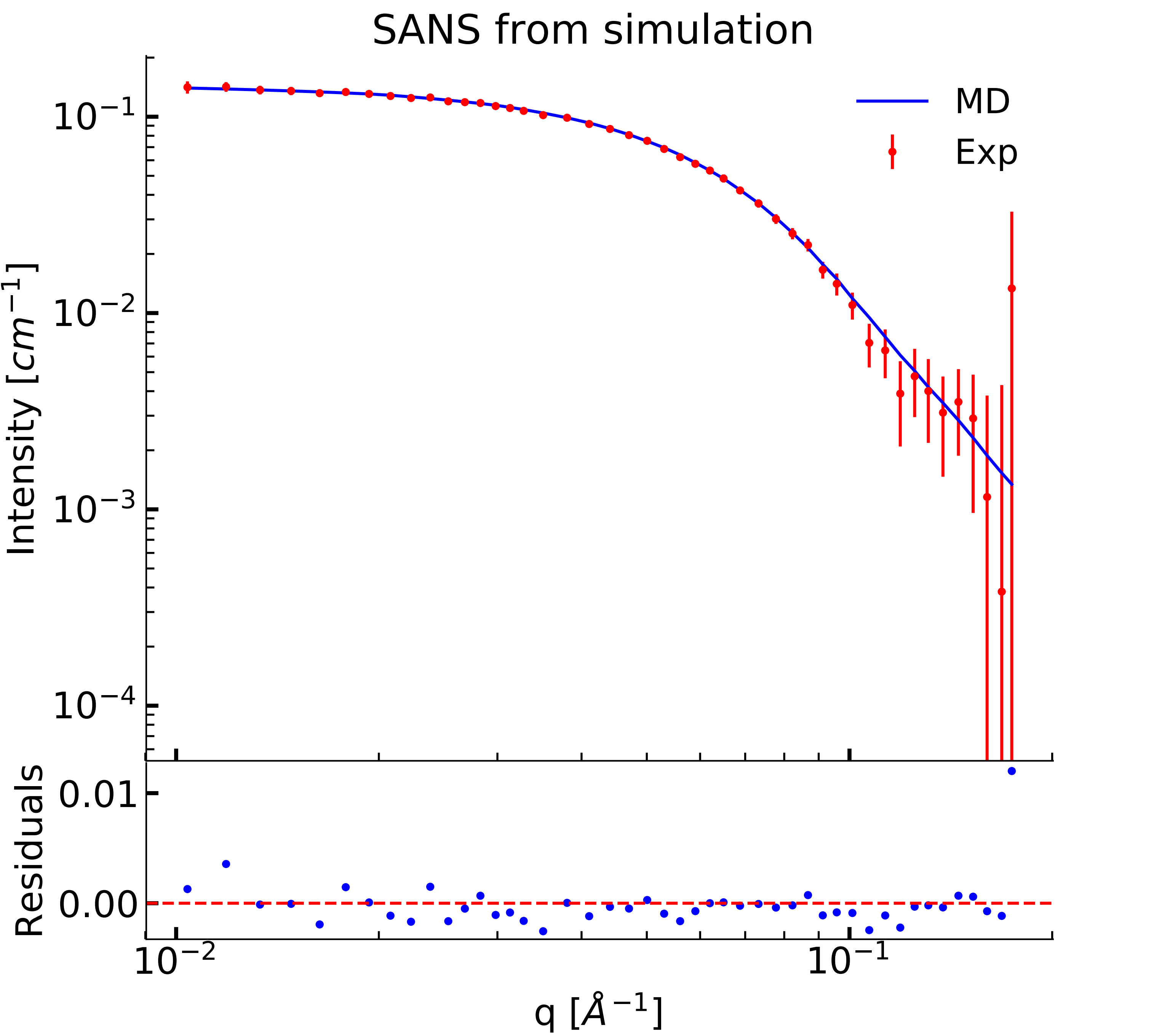
SANS from simulation
MD
Exp
Intensity [$cm^{-1}$]
$10^{-1}$
$10^{-2}$
$10^{-3}$
$10^{-4}$
Residuals
0.01
0.00
$10^{-2}$
$10^{-1}$
q [$Å^{-1}$]

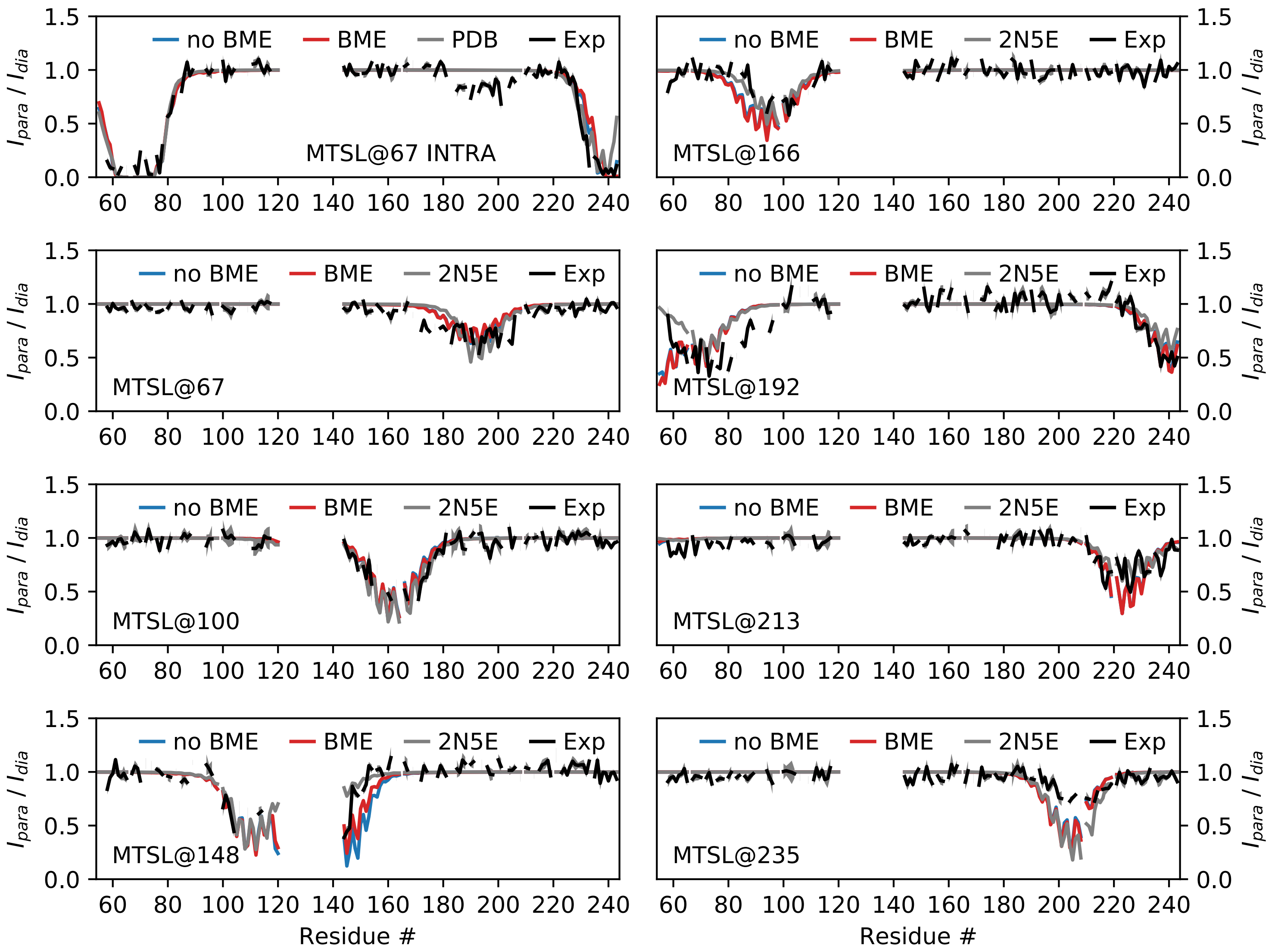

no BME
BME
PDB
Exp
MTSL@67 INTRA
2N5E
MTSL@166
MTSL@67
MTSL@192
MTSL@100
MTSL@213
MTSL@148
MTSL@235
$I_{para}$ / $I_{dia}$
Residue #

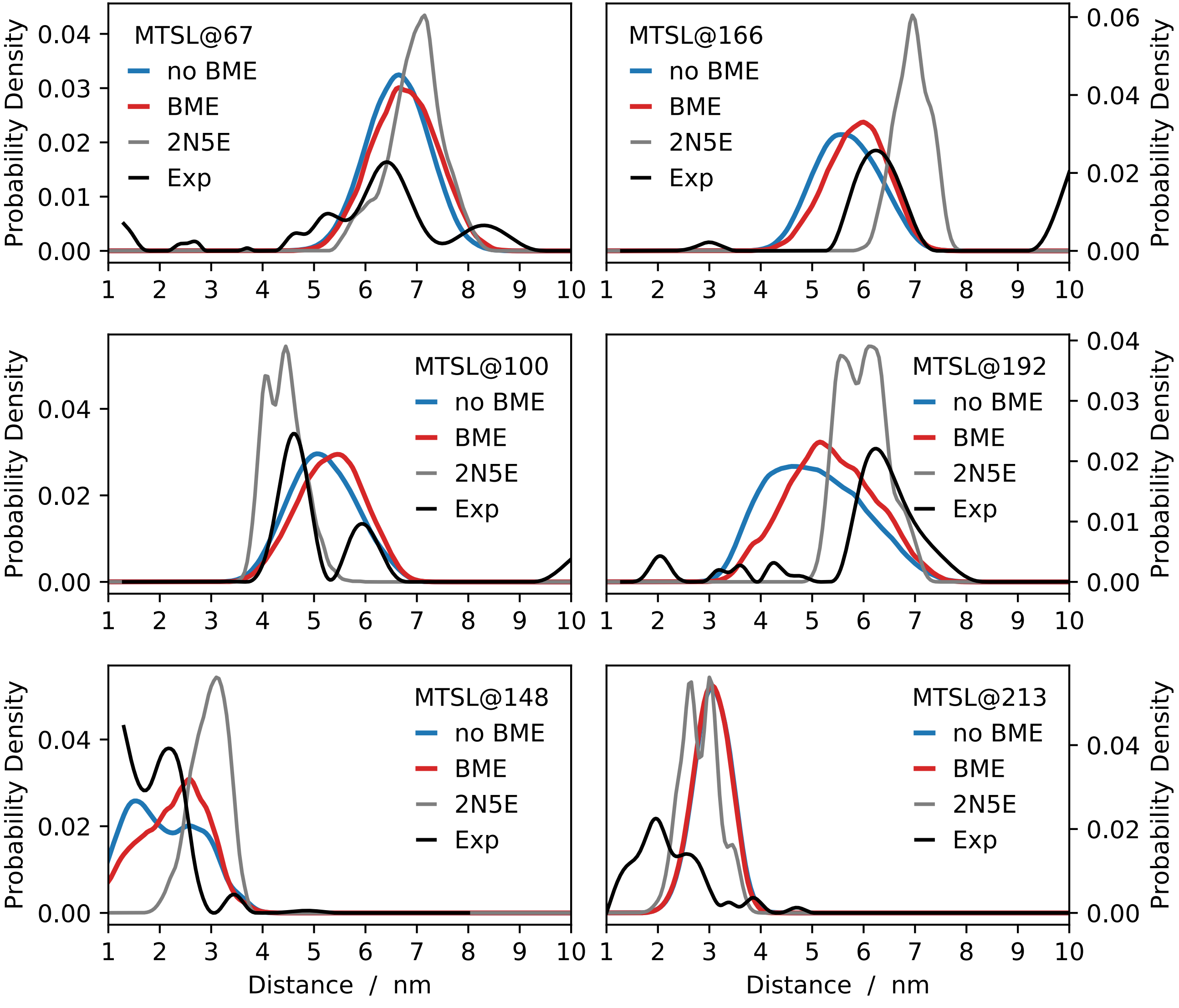

MTSL@67
MTSL@166
MTSL@100
MTSL@192
MTSL@148
MTSL@213
no BME
BME
2N5E
Exp
Probability Density
Distance / nm

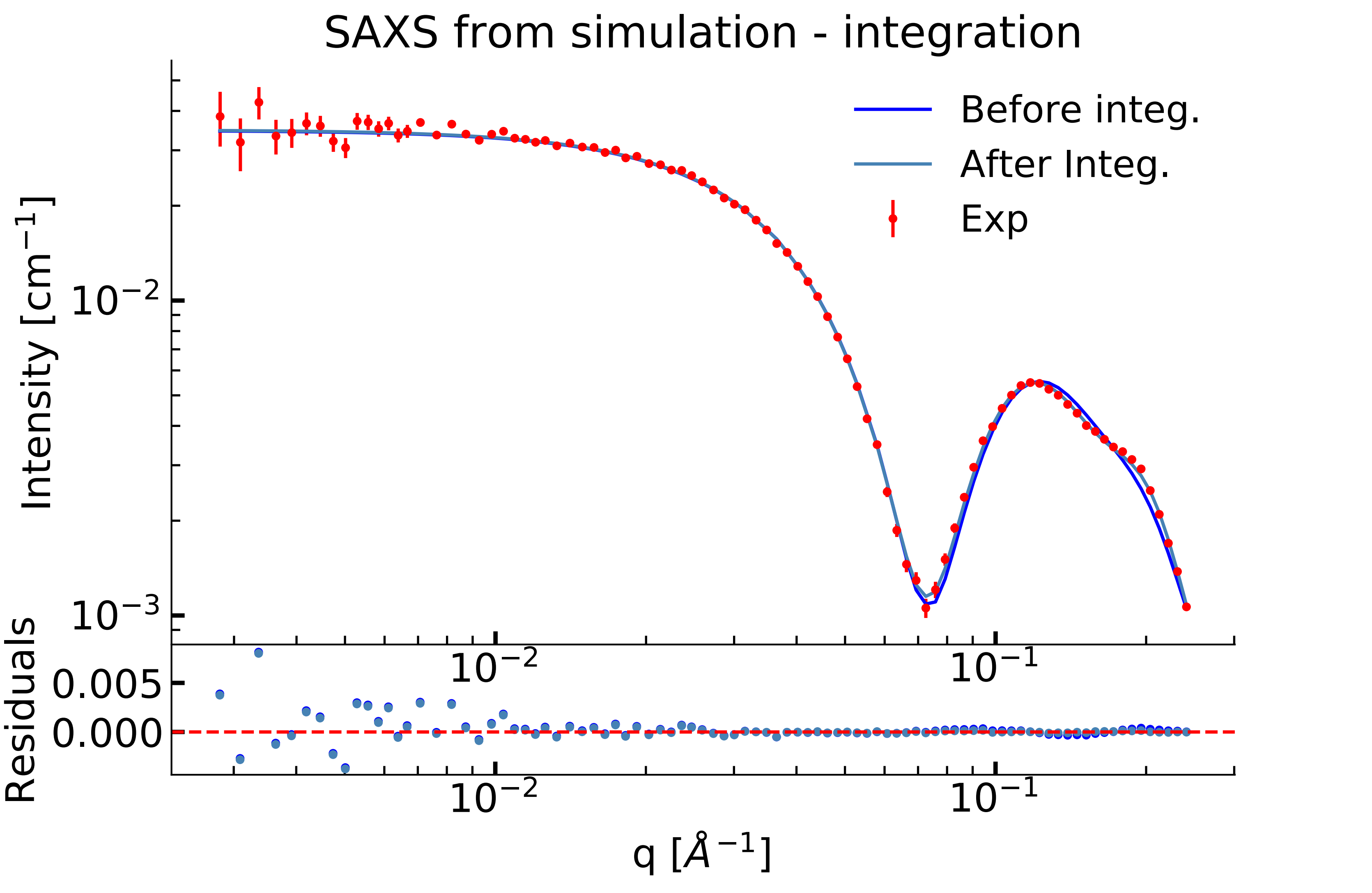

SAXS from simulation - integration
Before integ.
After Integ.
Exp
Intensity [cm$^{-1}$]
$10^{-2}$
$10^{-3}$
Residuals
0.005
0.000
$10^{-2}$
$10^{-1}$
q [$Å^{-1}$]

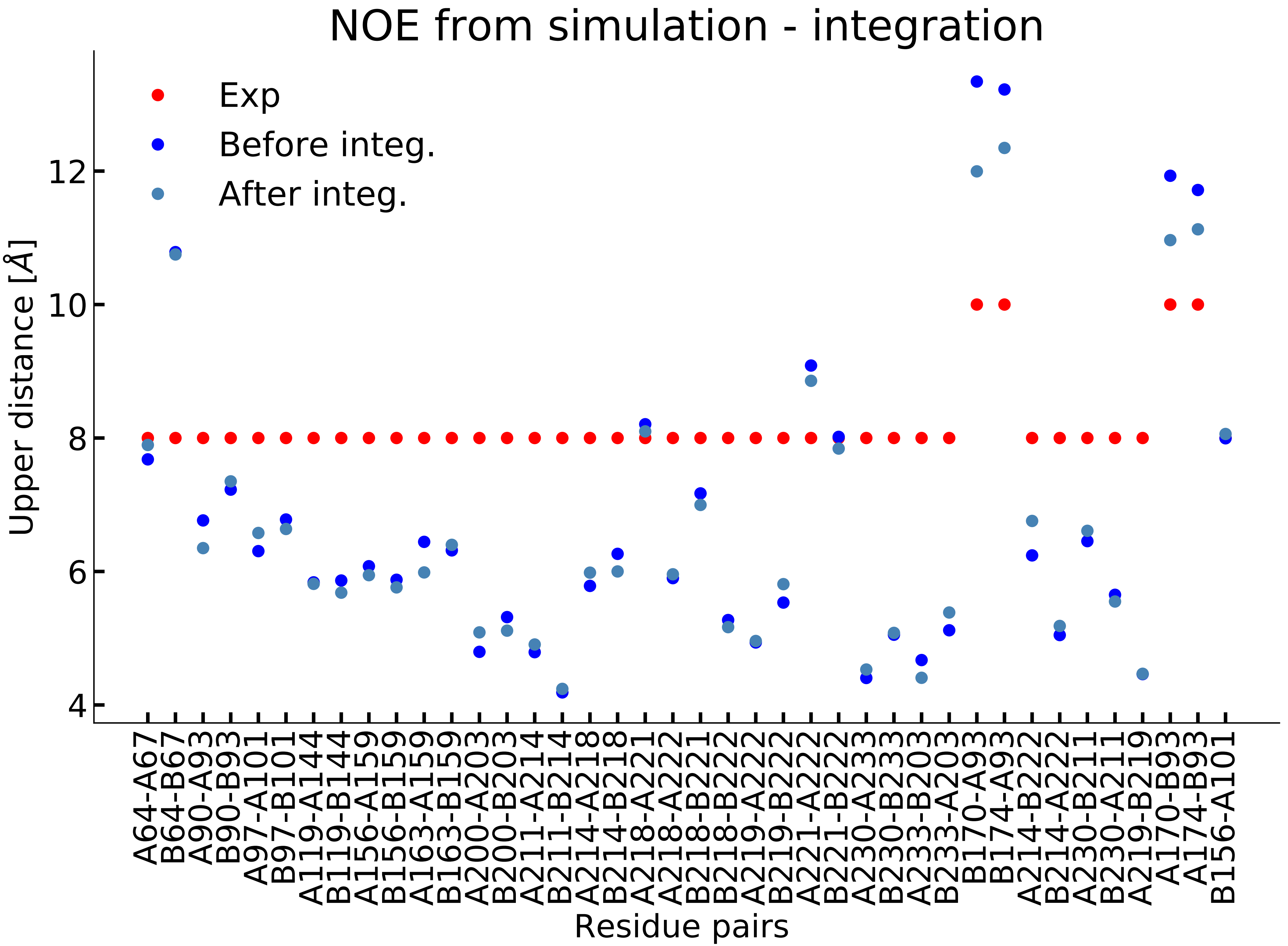

NOE from simulation - integration
Exp
Before integ.
After integ.
Upper distance [Å]
12
10
8
6
4
A64-A67
B64-B67
A90-A93
B90-B93
A97-A101
B97-B101
A119-A144
B119-B144
A156-A159
B156-B159
A163-A159
B163-B159
A200-A203
B201-B203
A211-A214
B211-B214
A214-A218
B214-B218
A218-A221
A218-A222
B218-B221
B218-B222
A219-A222
B219-B222
A221-A222
B221-B222
A230-A233
B230-B233
A233-B203
B233-A203
B170-A93
B174-A93
A214-B222
B214-A222
A230-B211
B230-A211
A219-B219
A170-B93
A174-B93
B156-A101
Residue pairs

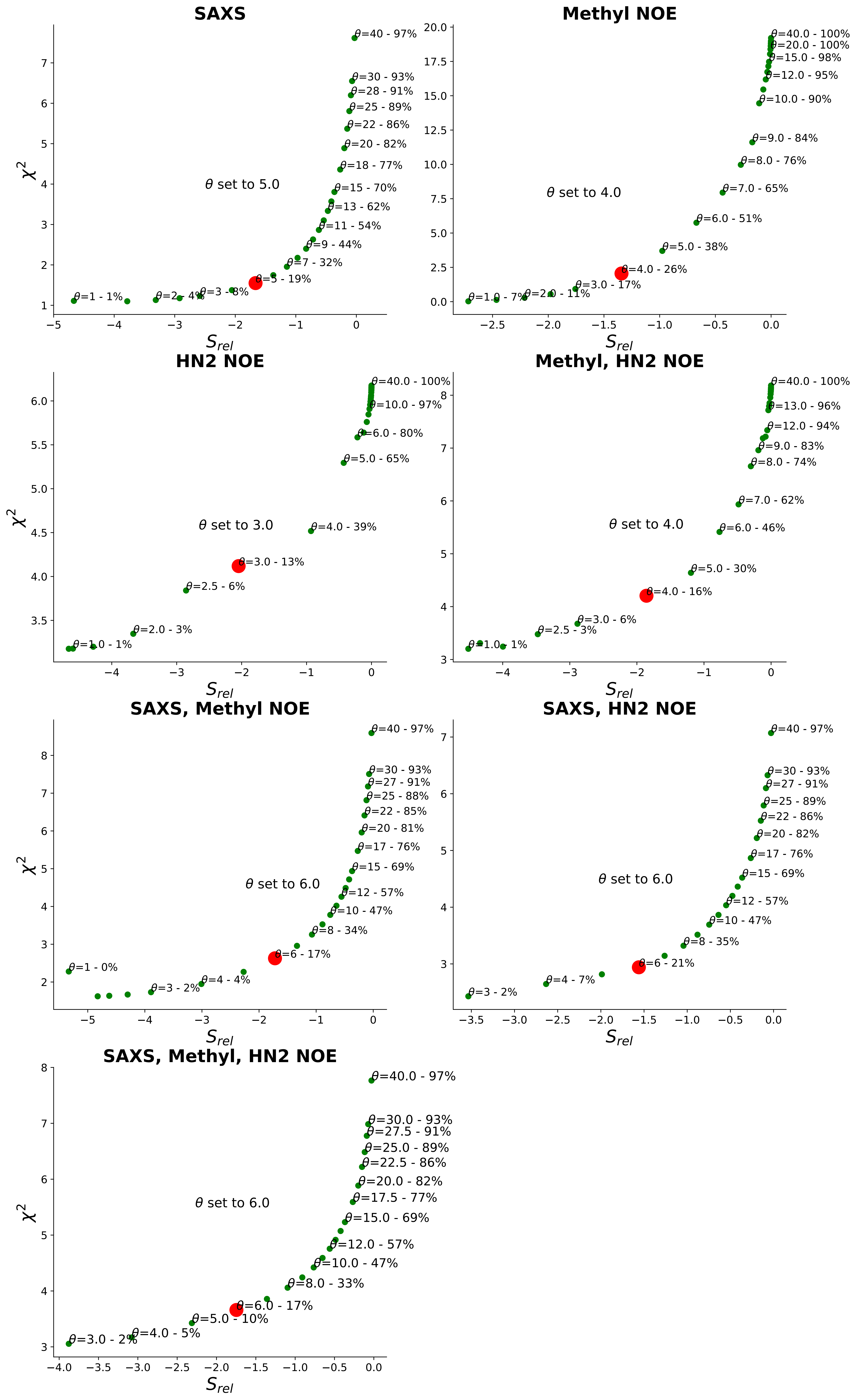
SAXS
θ set to 5.0
θ=1 - 1%
θ=2 - 4%
θ=3 - 8%
θ=5 - 19%
θ=7 - 32%
θ=9 - 44%
θ=11 - 54%
θ=13 - 62%
θ=15 - 70%
θ=18 - 77%
θ=20 - 82%
θ=22 - 86%
θ=25 - 89%
θ=28 - 91%
θ=30 - 93%
θ=40 - 97%
Methyl NOE
θ set to 4.0
θ=1.0 - 7%
θ=2.0 - 11%
θ=3.0 - 17%
θ=4.0 - 26%
θ=5.0 - 38%
θ=6.0 - 51%
θ=7.0 - 65%
θ=8.0 - 76%
θ=9.0 - 84%
θ=10.0 - 90%
θ=12.0 - 95%
θ=15.0 - 98%
θ=20.0 - 100%
θ=40.0 - 100%
HN2 NOE
θ set to 3.0
θ=1.0 - 1%
θ=2.0 - 3%
θ=2.5 - 6%
θ=3.0 - 13%
θ=4.0 - 39%
θ=5.0 - 65%
θ=6.0 - 80%
θ=10.0 - 97%
θ=40.0 - 100%
Methyl, HN2 NOE
θ set to 4.0
θ=1.0 - 1%
θ=2.5 - 3%
θ=3.0 - 6%
θ=4.0 - 16%
θ=5.0 - 30%
θ=6.0 - 46%
θ=7.0 - 62%
θ=8.0 - 74%
θ=9.0 - 83%
θ=12.0 - 94%
θ=13.0 - 96%
θ=40.0 - 100%
SAXS, Methyl NOE
θ set to 6.0
θ=1 - 0%
θ=3 - 2%
θ=4 - 4%
θ=6 - 17%
θ=8 - 34%
θ=10 - 47%
θ=12 - 57%
θ=15 - 69%
θ=17 - 76%
θ=20 - 81%
θ=22 - 85%
θ=25 - 88%
θ=27 - 91%
θ=30 - 93%
θ=40 - 97%
SAXS, HN2 NOE
θ set to 6.0
θ=3 - 2%
θ=4 - 7%
θ=6 - 21%
θ=8 - 35%
θ=10 - 47%
θ=12 - 57%
θ=15 - 69%
θ=17 - 76%
θ=20 - 82%
θ=22 - 86%
θ=25 - 89%
θ=27 - 91%
θ=30 - 93%
θ=40 - 97%
SAXS, Methyl, HN2 NOE
θ set to 6.0
θ=3.0 - 2%
θ=4.0 - 5%
θ=5.0 - 10%
θ=6.0 - 17%
θ=8.0 - 33%
θ=10.0 - 47%
θ=12.0 - 57%
θ=15.0 - 69%
θ=17.5 - 77%
θ=20.0 - 82%
θ=22.5 - 86%
θ=25.0 - 89%
θ=27.5 - 91%
θ=30.0 - 93%
θ=40.0 - 97%
χ2
Srel

| Data for integration | $\theta$ | $S_{rel}$ | $\chi^2$ | | |
|---|---|---|---|---|---|
| | | | SAXS | Methyl-NOE | HN-NOE |
| PDB | - | 0 | 2.9 | 20.7 | 6.3 |
| MD | - | 0 | 10.0 | 19.5 | 6.2 |
| MD + SAXS | 5 | -1.7 | 1.5 | 19.8 | 5.9 |
| MD + Methyl-NOE | 4 | -1.3 | 7.5 | 2.1 | 5.7 |
| MD + HN-NOE | 3 | -2.0 | 11.1 | 24.0 | 4.1 |
| MD + SAXS + Methyl-NOE | 6 | -1.7 | 1.9 | 8.2 | 6.3 |
| MD + SAXS + HN-NOE | 6 | -1.6 | 1.8 | 19.2 | 5.2 |
| MD + Methyl-NOE+ HN-NOE | 4 | -1.9 | 8.9 | 3.6 | 4.3 |
| MD + SAXS + Methyl-NOE + HN-NOE | 6 | -1.7 | 1.9 | 9.3 | 5.4 |
